# Similarities Between Bacterial GAD and Human GAD65: Implications in Gut Mediated Autoimmune Type 1 Diabetes

**DOI:** 10.1101/2021.11.29.470330

**Authors:** Suhana Bedi, Tiffany M. Richardson, Baofeng Jia, Hadeel Saab, Fiona S.L. Brinkman, Monica Westley

**Author notes:** authors contributed equally to this work (Suhana Bedi, Tiffany M. Richardson, Baofeng Jia). corresponding author, Corresponding author: Monica Westley Ph.D., the(sugar)science.

## Abstract

A variety of islet autoantibodies (AAbs) can predict and possibly dictate eventual type 1 diabetes (T1D) diagnosis. Upwards of 75% of those with T1D are positive for AAbs against glutamic acid decarboxylase (GAD65), a producer of gamma-aminobutyric acid (GABA) in human pancreatic beta cells. Interestingly, bacterial populations within the human gut also express GAD65 and produce GABA. Evidence suggests that dysbiosis of the microbiome may correlate with T1D pathogenesis and physiology. Therefore, autoimmune linkages between the gut microbiome and islets susceptible to autoimmune attack need to be further elucidated. Utilizing *silico* analyses, we show here that 25 GAD sequences from different human gut bacterial sources show sequence and motif similarities to human beta cell GAD65. Our motif analyses determined that a majority of gut GAD sequences contain the pyroxical dependent decarboxylase domain of human GAD65 which is important for its enzymatic activity. Additionally, we showed overlap with known human GAD65 T-cell receptor epitopes which may implicate the immune destruction of beta cells. Thus, we propose a physiological hypothesis in which changes in the gut microbiome in those with T1D result in a release of bacterial GAD, thus causing miseducation of the host immune system. Due to the notable similarities, we found between humans and bacterial GAD, these deputized immune cells may then go on to target human beta cells leading to the development of T1D.

## Introduction

Glutamic acid decarboxylase (GAD65) is a prominent autoantibody in type 1 diabetes (T1D). For those diagnosed as a teen, in addition to having an HLA DR3/DR2 signature, it is likely to be the first autoantibody clinically detected in the blood. GAD65 antibodies are prevalent in the majority of T1D cases, occurring in over 70% of patients [1]. Autoantibodies directed against GAD65 recognize linear and conformational epitopes throughout the protein. The functional region of GAD65 contains a catalytic group involving several residues that are distantly located in the primary amino acid sequence of the protein. The enzymatic function of GAD65 and subsequent regulation of GABA production is dependent on a key residue within the catalytic loop, Lys396 [2]. This residue covalently binds pyridoxal 5’-phosphate (PLP), a co-factor that is essential for the production of GABA by GAD65. Thus, T1D-related immune recognition of GAD65 is highly dependent on the exposure of these regions to autoreactive antigen-presenting cells (APCs). These activated APCs can then go on to attack GAD65-expressing pancreatic beta cells leading to their destruction and eventual T1D diagnosis.

In addition to beta cells, a multitude of bacteria within the gut express GAD (bacterial nomenclature for human GAD65). While there is great diversity within the gut microbiota, more than 95% of gut bacterial species fall into four major microbial phyla: Firmicutes, Bacteroidetes, Actinobacteria, and Protecteobacteria [3]. Bifidobacterium adolescentis, in particular, is an important gut bacterial GABA producer and has the highest prevalence of gad genes in their genomes [4]. These species provide a homeostatic environment that fosters and facilitates gut epithelial integrity and immunity, along with providing energy. Varied literature supports both a decrease in microbial diversity and an increase in a “leaky gut” phenotype throughout T1D [5].

GABA producing Bifidobacterium bacterial counts drop as the Firmicutes/Bacteroides (F/B) ratios decrease in T1D [6]. There is also a reduction in the abundance of Clostridium clusters IV and XIVa and mucin-degrading bacteria such as Prevotella and Akkermansia. An important metabolite produced by these symbiotic bacterial populations, butyrate, supports gut integrity and has beta cell-specific anti-inflammatory effects [7, 8, 9, 10, 11]. The reduction of these microbes and their byproducts may be a contributing factor to why children at risk for T1D have higher intestinal permeability and T1D-associated gut dysbiosis.

One hypothesis for the interrelationship between dysbiosis and T1D is that as these bacterial populations die, they release GAD65 mimetics that trigger the immune system. Presumably, as GABA-producing bacteria like B. adolescentis die, either from antibiotic misuse or other pathology, immune cells located in Peyer’s patches and deep in the enterocyte border detect the inappropriate presence of GAD. Thus, APCs can act to further inform CD8+ T cells in nearby lymph nodes. The lymphatic pathways between the intestinal submucosa and pancreatic lymph nodes may provide a route for these “miseducated” CD8+ T cells to gain access to GAD65+ beta cells. GAD65 may also originate from beta cells leading to an increase in epitope spreading and a misguided immune attack on these insulin-producing cells.

To initiate the exploration of this hypothesis, we utilized in silico methodologies to compare gut microbial species that decline at the onset of T1D and express GAD, with GAD65 sequences in pancreatic beta cells. We show that there is notable sequence similarity, including relevant motif and T-cell epitope overlaps between several bacterial species and beta cells particularly in the functionally important pyroxical dependent decarboxylase domain of GAD65. This in silico exercise suggests that further investigation is warranted of the relationship between microbial disappearance and appearance of GAD65 autoantibodies during the prodrome and presentation of T1D.

## Materials and Methods

### GAD Sequence and T cell receptor epitope curation

Twenty-five bacterial GAD proteins from species of interest, alongside those of mouse, chimpanzee, and human GAD65 sequences were curated from the NCBI genes database (Table 1). The bacterial species were chosen by considering their association with the gut microbiome, their significance in the gut microbiome alterations seen in T1D patients, human-related pathogen status, and non-human-related environmental species. GAD65 T cell receptor epitopes were curated from a literature review.

**Table 1.** GABA producing bacterial species and their respective GAD gene.

| Category | Species | Gene ID |
| --- | --- | --- |
| Human | Human | 2572 |
| Primate | Chimpanzee | 466026 |
| Mouse | Mouse | 14417 |
| T1D | *Streptococcus pneumoniae* | Z49109.1 |
| T1D | *Bifidobacterium moukalabense* | 61040350 |
| T1D | *Bifidobacterium adolescentis* | 56675515 |
| T1D | *Bacteroides fragilis* | 61595025 |
| T1D | *Ruminococcus* | Genome not found |
| T1D | *Veillonella* | Genome not found |
| T1D | *Blautia* | Genome not found |
| T1D | *Lachnospira* | Genome not found |
| T1D | *Alistipes obesi* | Genome not found |
| T1D | *Clostridium* spp. | 29570578 |
| T1D | *Prevotella* | Genome not found |
| T1D | *Parabacteroides distasonis* | CP040468 |
| Gut | *Escherichia coli* | 946058 |
| Gut | *Listeria monocytogenes* | 985123 |
| Gut | *Salmonella enterica* | 13913147 |
| Gut | *Streptomyces* sp. NRRL B-24085 | 6211654 |
| Gut | *Carnobacterium maltaromaticum* | 56848084 |
| Gut | *Nocardia brasiliensis* | 13733698 |
| Gut | Lactobacillaceae | 56991876 |
| Pathogen | Gamma proteobacteria | QNEL01000191.1 |
| Pathogen | *Klebsiella pneumoniae* IS22 | 12545741 |
| Environment | *Planctomycetes* bacterium | KAA3606119.1 |
| Environment | *Sediminibacterium* sp. | JADMJG010000006.1 |
| Environment | *Moorea producens* | CP017599.1 |
| Environment | Magnetococcales bacterium | JADGBX010000054.1 |
| Environment | *Scytonema hofmannii* | ANNX02000047.1 |
| Environment | Candidatus Aminicenantes bacterium | WVZL01000014.1 |
| Environment | *Bdellovibrio* sp. qaytius | CP025734.1 |
| Environment | Verrucomicrobia | NZEQ01000004.1 |

All GAD sequences in bacteria and the three mammalian species under consideration were obtained by performing a BLASTP search, with human GAD65 as the query sequence. Furthermore, only those sequences with significant e and p values were well annotated on the NCBI database and/or RefSeq and were identified as GAD proteins were considered. All other hits, including unnamed protein products, were ignored.

### Multiple Sequence Alignment and Motif Identification

Multiple sequence alignment was performed using ClustalW (Ver. 1.2.2) with a gap opening penalty of 10 and a gap extension penalty of 0.2 [12]. The resulting alignment is visualized and annotated with CLC Sequence Viewer (Ver. 8.0) to highlight conservation and amino acid properties. Identification of ungapped motifs was done using the MEME tool (Ver. 5.4.1) under the MEME suite tools for motif discovery and enrichment [13]. The output consisted of 10 motifs, shared by various (not necessarily all) organisms into consideration. Subsequently, each motif was run against protein families, using a Pfam (Ver 33.1) search. The Motifs identified by MEME were mapped against the query sequences using MAST(Ver. 5.4.1), to obtain the coordinates of motifs, in each gad sequence [13].

### Logo plot & T Cell Receptor Epitopes

The multiple sequence alignment obtained was converted into a logo plot, using Weblogo (Ver 2.8.2) [14]. Here, each logo consists of a stack of symbols, and each symbol’s height is indicative of the amino acid conservation, at a particular location. All motifs and T-cell receptor epitope coordinates, obtained in the previous steps, were marked on the logo plot. The plot was then trimmed, with a special emphasis on regions with high overlap between motifs and T-cell receptor epitopes.

## Results

### Multiple sequence alignment highlights several conserved residues between human and bacterial GAD

A total of 25 GAD protein sequences, including 3 mammalian GAD65 (human, chimpanzee, and mouse) and 22 bacterial GAD, were aligned (Table 1 & Fig. 1). The 3 mammalian GAD65 exhibited high conservation with each other, while 20 out of the 22 bacteria GAD exhibited high similarity to one another. Two bacteria, *Bdellobibrio sp. Qaytius* and *Sediminibacterium spp.* did not exhibit any protein conservation. This may be due to a misannotation within the RefSeq genome. We also found interesting similarities and differences between the mammalian and bacterial GAD sequences. There were two regions of interest: AA389-397 and AA418-425. AA389-397 contained several charged residues interspaced with hydrophobic residues, potentially a conserved alpha-helix structure. AA418-AA425 contains conserved HK residues. These two residues represent a known pyridoxal phosphate-binding site in *E.coli* GAD (H275 and K276), that is likely conserved in mammalian GAD65. The length of mammalian GAD65 was ~120 amino acids longer than bacterial GAD. Based on our alignment, this difference is mostly centered around the N-terminus, which suggests adaptations and inclusions of signal peptides required for complex cellular trafficking in mammalian cells. Also, we identified a few small gaps within the core regions of GAD between species, these gaps are unlikely of any biological significance.

**Figure 1.**
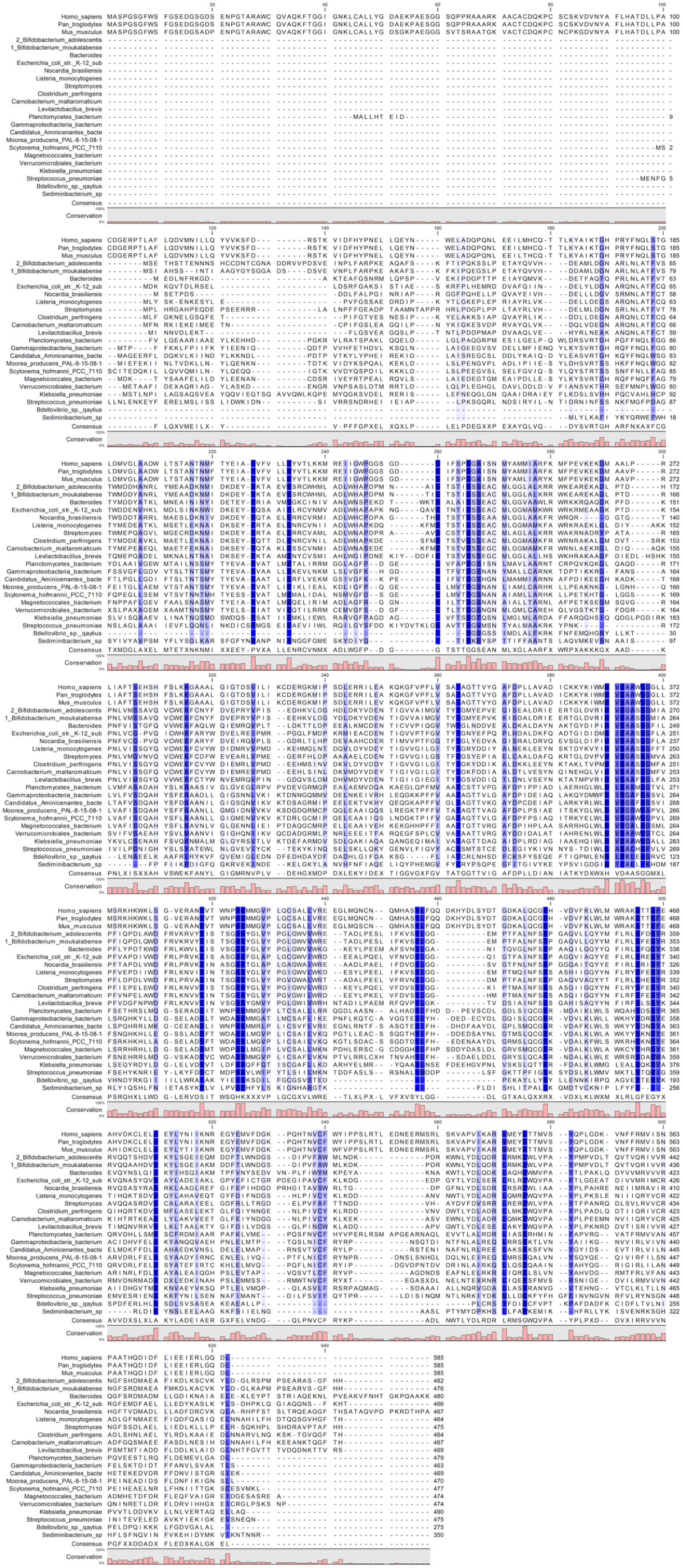
Multiple sequence alignment of twenty bacterial GAD protein and three animal GAD65 protein highlights conserved residues. Across a diverse number of bacterial species and 3 animal species, 12 amino acids were found to be 100% conserved (shaded in blue). Amino acids 1-110 are likely signal peptides required for eukaryotic protein translation and cellular transport not needed in bacteria. H275 and K276 are key residues required for PDP binding and are conserved in human GAD65 and bacterial GAD.

### Motif analysis uncovers regional similarities of human and bacterial GAD enzymatic regions

Our motif analysis found 10 ungapped motifs with each motif present in at least 10 of the sequences under consideration (Fig. 2). Even though multiple common motifs between human GAD65 and bacterial GAD can be observed, it is noteworthy that two motif blocks are conserved all across the organisms considered, which is indicative of the sequence similarity observed in the multiple sequence alignment. In addition, except for *Streptococcus pneumoniae*, one motif block was consistently present in all sequences. Furthermore, these motifs were queried on the Pfam and the CDD databases, and all 10 motifs were identified as matches to the pyridoxal-dependent decarboxylase (PDP) conserved domain, which is instrumental in the biosynthesis of GABA, from glutamate. This indicates that human GAD65 and bacterial GAD share sequence similarities in a functional region of GAD.

**Figure 2.**
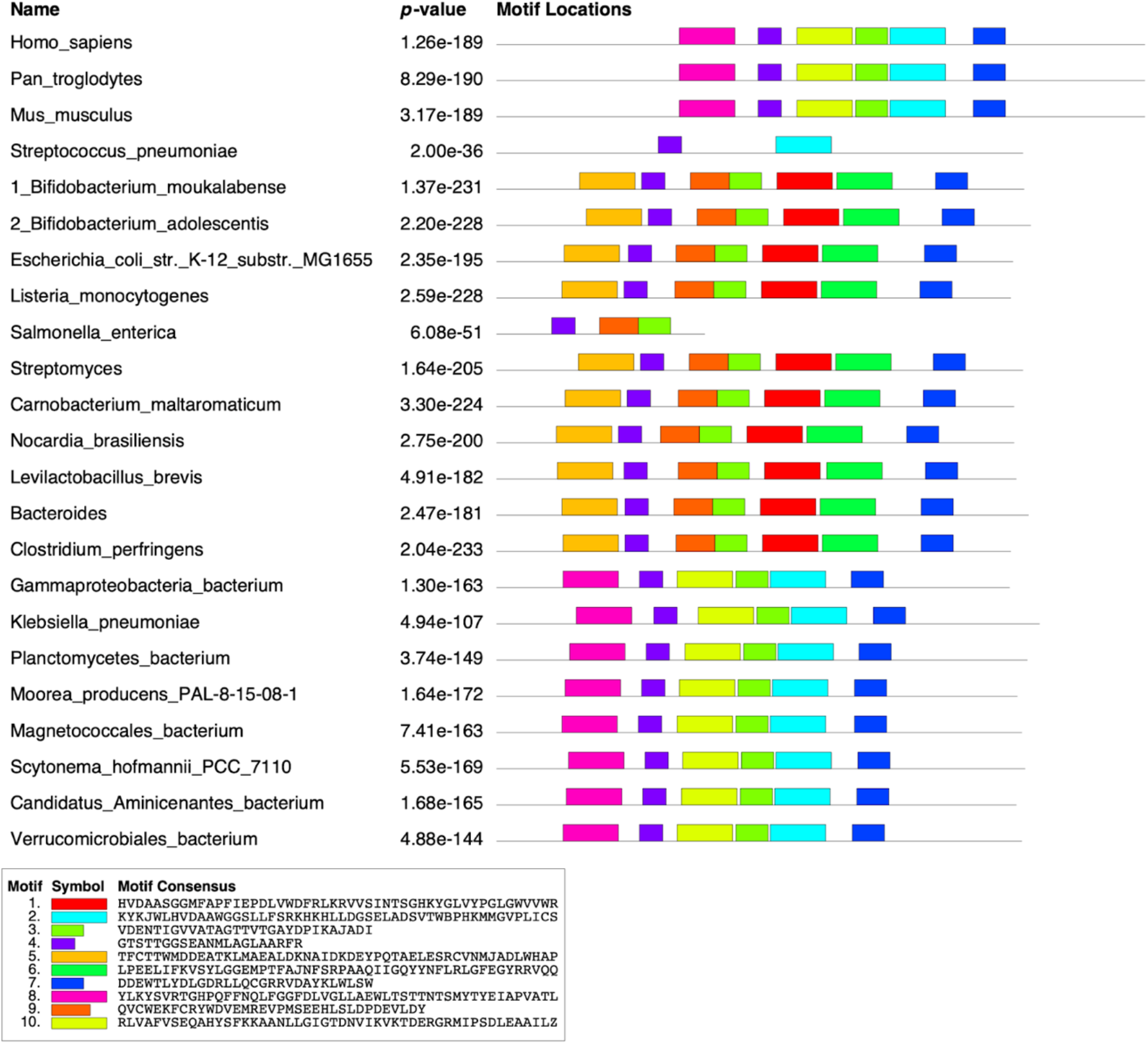
Motif Discovery Analysis of twenty bacterial GAD protein and three animal GAD65 proteins to identify motifs conserved across various species and their functional relevance with GAD protein. Across all 23 GAD sequences under consideration, 10 motifs were each found in at least 10 of the sequences used for motif discovery. Each motif appears in a separate color below with the individual sequences described as well. In addition, 2 of the 10 motifs were found in all of the sequences. Each motif was then queried for, using Pfam and CDD searches and it was found that all 10 of the motifs, hit the PDP domain in the GAD65 protein. All motifs match the pyridoxal-dependent decarboxylase (PDP) conserved domain.

### Overlap exists between T cell receptor epitopes of GAD65 and bacterial GAD

To compare the overlap between T cell receptor epitopes, human GAD65, and bacterial GAD, a logo plot of the multiple sequence alignment was made using three representative bacterial GAD and three animal GAD65 proteins. Motif positions for each sequence were manually annotated using the motif coordinates obtained in motif discovery and enrichment analysis. Thereafter, the logo plot was annotated with T cell receptor epitopes and the overlapping regions were observed. The fine lines indicate the motif coordinates, while the red blocks indicate the CD8+ epitopes and blue blocks indicate the CD4+ epitopes. Three regions overlapped with a motif, which as stated in the motif analysis result description, hit the PDP domain (Fig. 3). This was also confirmed by a CDD search. Of the amino acids in which both CD4 and CD8 receptor epitopes overlap, there is at least one amino acid residue that is conserved across bacterial GAD and GAD65 (Fig. 4). Such overlaps emphasize a possible explanation for how T cells could recognize both bacterial and human GAD65.

**Figure 3.**
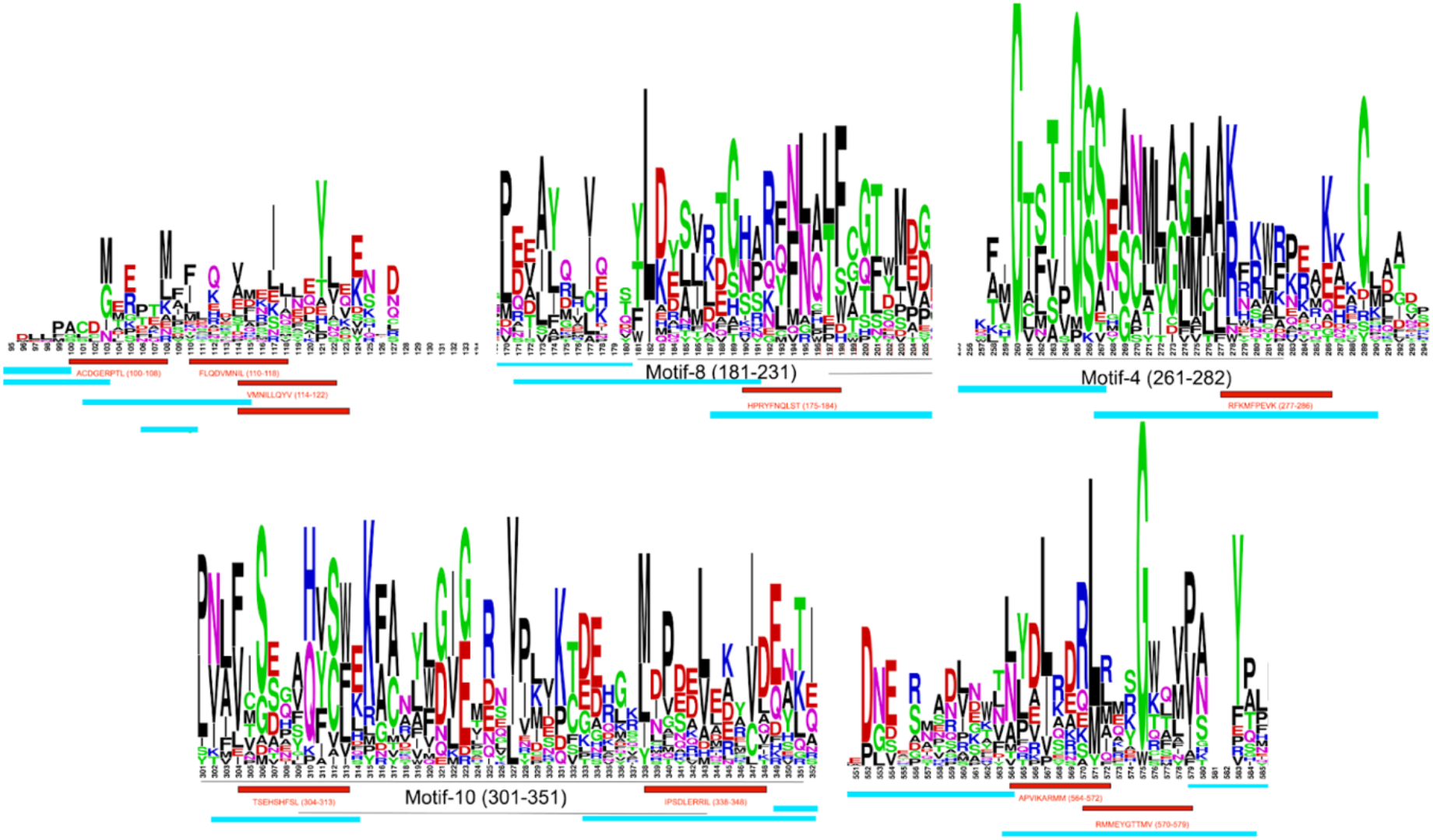
Conserved regions of GAD share common CD8+ and CD4+ T cell receptor epitopes. A total of five regions of interest were identified by overlapping T cell receptor epitopes to the sequence logo of three representative bacterial GAD and three animal GAD65 proteins. Three regions contained a motif match to Group II pyridoxal-dependent decarboxylases. Of the amino acids in which both CD4+ and CD8+ receptor epitopes overlap, there is at least 1 amino acid residue that is conserved across bacterial GAD and GAD65. Red is CD8+ epitopes and Blue is CD4+ epitopes.

**Figure 4.**
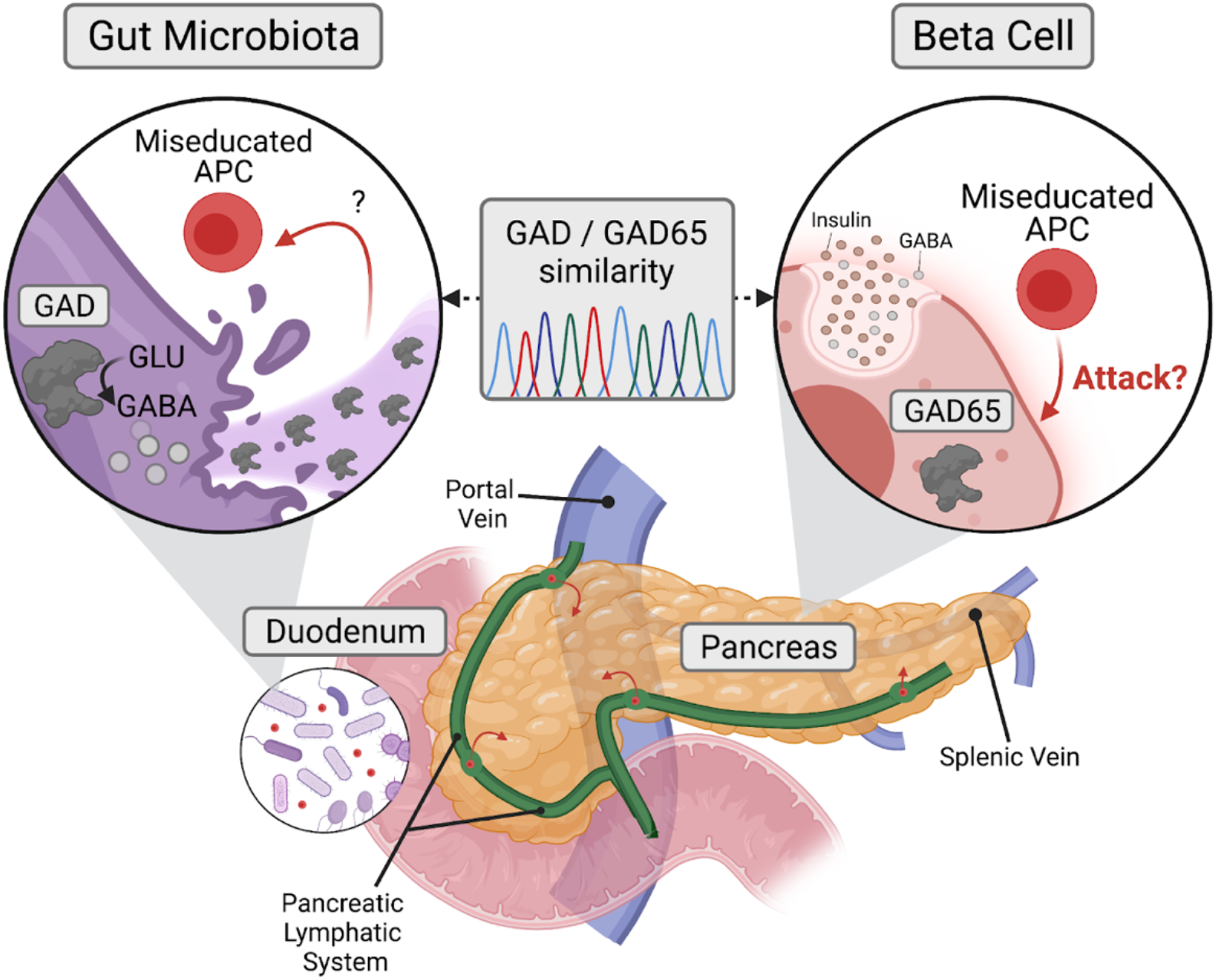
Miseducation of antigen-presenting cells (APCs) by bacterial GAD may contribute to pancreatic beta cell immune-mediated destruction. The death of GABA-producing bacteria at the onset of T1D may release bacterial GAD65 which could lead to the miseducation of APCs. Therefore, this miseducation could provoke the immune system to attack and produce antibodies targeting human GAD65 expressing beta cells because of the similarity and conservation between bacterial GAD and human GAD65. The pancreatic lymphatic system could provide a pathway for these miseducated APCs to traverse from the duodenum into pancreatic islets.

## Discussion

In this study, we explored the sequence similarities between GAD in twenty GABA-producing bacteria and GAD65 in humans. We found that human and bacterial GAD have similar motifs and conserved residues. We identified notable conservation centered around the pyridoxal dependent decarboxylase domain of GAD65 found in pancreatic beta cells, particularly around two key substrate-binding residues. Moreover, some conserved sequences in bacterial GAD overlap with known human GAD65 T cell receptor epitopes. As one example, GAD65-specific HLA-DR4 (DRB1*0401)–restricted murine T-cell hybridoma line T33.1 recognizes the GAD65 274-286 epitope, and this epitope was conserved across bacteria and human samples examined here (Figure 1). This *in silico* proof-of-concept suggests a possible relationship between disappearance of GAD containing GABA-producing bacteria in the human gut microbiome and the appearance of GAD65 autoantibodies during the onset of T1D that needs to be further elucidated.

It is important to note a few limitations within our analyses. Many gut microbial genomes and genome annotations were incomplete, particularly for some GABA-producing bacterial species of importance that are significantly changed in T1D. We did not include several species due to missing GAD genome annotations. Together, only 20 sequences were analyzed for their homology, with many belonging to environmental organisms. A larger cohort would further strengthen our sequence analyses and hypotheses. Furthermore, we hypothesized that the change in anti-GAD65 is partly due to the changes in the gut microbiome. Additional studies are needed to determine what insult(s) would cause such a disturbance and what cellular mechanisms are involved in the miseducation of immune cells due to the release of GAD in the gut. The window for the disappearance of the *gad* containing bacterial species should be identified as to whether it precedes or follows the appearance of autoantibodies to GAD65 in humans. Lastly, the analysis relied on known and curated T cell epitopes of GAD65 from the literature review, in which we demonstrated similarity to bacterial GAD. Whether or not the bacterial peptides bind to human T cell receptors needs to be further verified *in vitro.*

We postulate that bacterial GAD found in GABA producing bacteria may act as an antigen to activate submucosal T cells as a result of gut microbiome bacterial destruction, either viral or antibiotic induced. Further, we posit these primed bacterial GAD T cells bind GAD65 in the islet environment due to the GAD and GAD65 epitope similarity. That leaves us to explore how GAD65, an intracellular enzyme found in alpha and beta islet cells, might get recognized by bacterial derived anti-GAD T cells.

In beta cells, GAD65 is intracellular and interacts with the cytosolic leaflet of Golgi membranes and secretory vesicles due to its post-translational palmitoylation (lipidation) at cysteines 30 and 45 [15]. GAD65’s shuttling to the Golgi for palmitoylation requires a signal located between residues 1-27 of the protein [16, 17]. GAD65 may transiently interact with the cell’s cytosolic leaflet upon exocytosis of beta cell granules [18]. Pre-clinical studies demonstrated that GAD65 can associate with the plasma membrane and enter the extracellular space [19].

GAD65 is certainly present in alpha and beta cells and may interact with a GAD T cell from the gut at the plasma membrane of both islet cell types, but there may be another route of contact. A recent review highlights how beta cells can become “deputized” and act as antigen-presenting cells (APCs) under certain conditions [20]. In this scenario, “deputized” beta cells that present GAD65 might have an opportunity to interact with a GAD sensitized T cell. Endothelial cells in the highly vascularized islet environment have been named as additional “deputized” APCs. These endothelial cells may be exposed to transient plasma membrane resident GAD65, or GAD65 containing EVs from stressed pancreatic beta cells [21, 22] and act to present them to anti-GAD gut derived T cells.

A “two-hit” scenario is one where bacterial GAD primed T cells recognize alpha and beta cell derived GAD65 to impact both intra islet cell interaction as well as simultaneously disrupting local immune response. In islet α-cells, GABA induces membrane hyperpolarization and suppresses glucagon secretion, while in islet β-cells GABA induces membrane depolarization and increases insulin secretion [23]. One consequence of a decrease of GAD65 in the islet environment is the resultant decrease in GABA production resulting in hyperglycemia which is toxic to beta cells. GABA is also an immune cell modulator, shown to increase Tregs, decrease Th1 and CTLs fostering a tolerant immune environment [23]. A GAD65 depletion leading to a GABA decrease in the islet could pack a double punch of generating dysfunctional alpha and beta cells operating in a more highly reactive immune micro-environment.

One can imagine a scenario where GAD primed T cells from the gut, become initially alerted to the islet environment as GAD65 is presented via deputized alpha, beta, endothelial cells or some combination of the three. Once alerted, GAD primed T cells destroy beta or alpha cells that display GAD65 at the plasma membrane, reducing the overall GABA levels in the islets with resulting consequences of hyperglycemia which destabilizes beta cells, in addition to altered immune phenotype that favors attack of the islets.

Taken together, our *in silico* analyses of genome sequences and motifs from bacterial, human, and T cell epitopes identified several homologies that may inform further experimentation and inquiry into the relationship between T1D pathogenesis and microbiome dysbiosis.

## Acknowledgments

We wish to express our gratitude to the entire volunteer team at the(sugar)science and to our donors and sponsors that help us continue our mission of helping scientists who study Type 1 diabetes connect, collaborate, and accelerate research.

## Funding

the(sugar)science is an all-volunteer 501c3 non-profit organization. B.J. was supported by a Canadian Institutes of Health Research (CIHR) doctoral scholarship. T.M.R. was supported by a National Science Foundation Graduate Research Fellowship, and F.S.L.B. has a Simon Fraser University distinguished professorship and support from Natural Sciences and Engineering Research Council of Canada (NSERC).

## Author Contributions

M.W. devised the original experimentation and acquired funding; M.W., S.B., and B.J. contributed to the conceptualization; S.B. and B.J. performed the investigation and formal analysis; M.W., F.S.L.B, S.B., B.J., and T.M.R. interpreted the results; M.W., T.M.R., B.J., S.B., and H.S. wrote the original draft; all authors reviewed and edited the manuscript.

## Notes

### Competing Interest Statement

The authors have declared no competing interest.

### Summary of Updates

We are sending this pdf with corrected versions of the figures and their associated descriptions.

